# Reassessing the Functional Significance of BOLD Variability

**DOI:** 10.1101/2023.02.06.527384

**Authors:** R.P. Roberts, K. Wiebels, D. Moreau, D.R Addis

## Abstract

BOLD variability (SD_BOLD_) has emerged as a unique measure of the adaptive properties of neural systems that facilitate fast, stable responding, based on claims that SD_BOLD_ is independent of mean BOLD signal (mean_BOLD_) and a powerful predictor of behavioural performance. We challenge these two claims. First, the apparent independence of SD_BOLD_ and mean_BOLD_ may reflect the presence of deactivations; we hypothesize that while SD_BOLD_ may not be related to raw mean_BOLD_ it will be linearly related to *absolute* mean_BOLD_. Second, the observed relationship between SD_BOLD_ and performance may be an artifact of using fixed-length trials longer than response times. Such designs provide opportunities to toggle between on- and off-task states, and fast responders likely engage in more frequent state-switching, thereby artificially elevating SD_BOLD_. We hypothesize that SD_BOLD_ will be higher and more strongly related to performance when using such fixed-length trials relative to self-paced trials that terminate upon a response. We test these two hypotheses in an fMRI study using blocks of fixed-length or self-paced trials. Results confirmed both hypotheses: (1) SD_BOLD_ was robustly related with *absolute* mean_BOLD_; and (2) toggling between on- and off-task states during fixed-length trials reliably contributed to SD_BOLD_. Together, these findings suggest that a reappraisal of the functional significance of SD_BOLD_ as a unique marker of cognitive performance is warranted.

## Introduction

Over the last decade, a growing body of evidence has implicated intra-individual variability in the brain’s blood oxygen level dependent (BOLD) response—the signal measured by functional magnetic resonance imaging (fMRI)—as a unique and informative metric of brain function and cognitive capacity (Boylan et al., 2020; Burzynska et al., 2015; Garrett et al., 2010; Garrett, Kovacevic, et al., 2011; Garrett, McIntosh, et al., 2011; Garrett, Kovacevic, et al., 2013; Garrett et al., 2014; Gaut et al., 2019; Grady & Garrett, 2018; Guitart-Masip et al., 2016; Protzner et al., 2013). This perspective on BOLD variability (SD_BOLD_) rests on two separate claims. First, SD_BOLD_ is hypothesised to capture important properties of brain dynamics associated with aging and cognitive performance, with a body of research showing that BOLD variability tends to systematically decrease across the lifespan in many cortical regions (e.g., Garrett et al., 2010, 2013; Garrett, Kovacevic, et al., 2011; Grady & Garrett, 2018, but see Boylan et al., 2020) and that increased variability is associated with increased cognitive performance (Garrett, Kovacevic, et al., 2011, 2013; Garrett, Samanez-Larkin, et al., 2013). Importantly, in these studies, the relationships of both age and cognitive performance with task-related BOLD variability are stronger than those evident with traditionally-used measures of average BOLD signal (mean_BOLD_), indicating that SD_BOLD_ is a promising and informative measure of brain function. Second, it has been proposed that SD_BOLD_ is a measure of the BOLD signal that is independent from mean_BOLD_, thus providing unique yet complementary insights into brain function (Garrett et al., 2010, 2014).

With respect to the relationship between BOLD variability (and neural variability more generally) and cognitive performance, a number of hypotheses have been advanced (for a review, see Garrett, Samanez-Larkin, et al., 2013). For example, it has been suggested that variability indexes the dynamic range of neural systems; that is, brains exhibiting greater variability are capable of occupying a greater range of states, making them capable of responding more efficiently to dynamically changing environments (Grady & Garrett, 2014). Alternatively, BOLD variability may reflect a level of background neural noise that allows for stochastic resonance—the notion being that an optimal amount of internal noise leads to better detection of weak signals (McDonnell & Ward, 2011). Regardless of the precise mechanisms involved, the common interpretation of the relationship between BOLD variability and cognitive performance is that SD_BOLD_ captures a facilitative, “trait-like” property of brain function that plays a causal role in how individuals perform in a range of cognitive tasks (Waschke et al., 2021).

An alternative—and as yet, untested—explanation for the relationship between task-based SD_BOLD_ and cognitive performance is that it is an artefact of experimental designs used in previous studies. In these studies, cognitive performance is almost universally based on response times in blocked fMRI designs (i.e., better cognitive performance is measured by faster, more stable responses; e.g., Garrett, Kovacevic, et al., 2011, 2013; Garrett, Samanez-Larkin, et al., 2013; Grady & Garrett, 2018). Furthermore, the length of trials within the task blocks are fixed, and these fixed durations are substantially longer than the vast majority of response times.

Critically, this feature of the design provides opportunities for participants to toggle between on- and off-task states: they are engaged in the task for a short period of the trial and then, following their response, disengage prior to the onset of the next trial. If this toggling between on- and off-task states results in fluctuations in the BOLD signal, and if it occurs more markedly for individuals who consistently respond quickly, then fast responders (with low response time variability) would exhibit greater BOLD variability. Importantly, if this is true, it would undermine the argument that high BOLD variability captures a trait of neural systems that facilitates better cognitive performance. Rather, better performance (i.e., fast and stable response times) induces variability in the BOLD signal that is directly attributable to the specifics of the experimental design.

In terms of the relationship between SD_BOLD_ and mean_BOLD_, the notion that mean_BOLD_ and SD_BOLD_ provide unique brain metrics is based primarily on two findings. Examination of spatial maps showing voxels in which SD_BOLD_ is reliably related to age and/or cognitive performance show minimal spatial overlap with voxels in which mean_BOLD_ shows relationships with age/performance (e.g., Garrett, Kovacevic, et al., 2011). In addition, directly assessing the voxelwise linear relationship between the two measures of the BOLD signal yields very small effect sizes (*r* = −.02, Garrett, Kovacevic, et al., 2011; r = .04, Garrett et al., 2014). These findings suggest that information about mean_BOLD_ effects is largely uninformative in predicting SDBOLD.

However, assessing the voxelwise linear relationship between the two measures of the BOLD signal across all voxels in the brain may not be a valid measure of independence as it is potentially confounded by the presence of mean_BOLD_ deactivations. That is, it could be the case that the set of voxels that systematically activate in response to a task (high, positive mean_BOLD_) as well as the set that systematically decrease in BOLD signal (low, negative mean_BOLD_) exhibit increased SD_BOLD_, producing a U-shaped, quadratic relationship across all voxels between the two measures that is not captured by testing for a linear relationship.

In the current pre-registered study, we critically assess these two claims made of BOLD variability: i) that SD_BOLD_ plays an instrumental role in facilitating fast, stable responding in cognitive tasks; and ii) SD_BOLD_ is independent of mean signal. First, to determine whether the relationship between BOLD variability and response times is a direct result of experimental design, we compared SD_BOLD_ during blocks of fixed-length trial (*start-stop* design) with SD_BOLD_ during blocks of self-paced trials which, upon a response, terminate and immediately proceed to the next trial (*continuous* design). We predicted that if state-switching (i.e., the toggling between on- and off-task states) contributes to BOLD variability, then SD_BOLD_ should be reliably increased in start-stop relative to continuous designs. Furthermore, we predicted that SD_BOLD_ should be more strongly related to response times in the start-stop design relative to the continuous design. To test the second claim, we performed voxelwise correlations between mean_BOLD_ and SD_BOLD_ in a similar vein to previous research (Garrett et al., 2014), and also between *absolute* mean_BOLD_ and SD_BOLD_, and predicted that only the latter set of correlations should produce reliable relationships as it will be sensitive to the relationship between SD_BOLD_ and both mean_BOLD_ activations and deactivations.

## Methods

The study was pre-registered on the Open Science Framework and can be accessed at https://osf.io/svmjn. All analyses described below are clearly labelled as either confirmatory (i.e., specified in the pre-registration) or exploratory.

### Participants

23 right-handed young adults (aged 21–35 years, *M* = 26.87, *SD* = 3.89, 8 males) participated in this study, and provided informed consent in a manner approved by The University of Auckland Human Participants Ethics Committee. All participants were fluent in English, and had no history of neurological and psychiatric disorders. Participants were compensated for their time with a gift certificate valued at $25 NZD.

### Materials and Procedure

#### Cognitive Tasks

Participants performed a visual task and a working memory task during fMRI scanning (see Figure 1); both were based closely on two tasks used in previous work on BOLD variability (e.g., Garrett, Kovacevic, et al., 2011). The visual task was a perceptual matching task, in which participants were presented with a target Gabor patch in the middle of the upper portion of the screen, and three Gabor patches in the bottom half of the screen, and had to indicate which of the bottom three patches matched the orientation of the target. The working memory task was a delayed match-to-sample task, in which a target Gabor patch was presented by itself on the screen for 1500ms (initial display), followed by blank screen (delay interval) for 2500ms, and then a test display of three Gabor patches presented in the lower half of the screen. Participants were required to select which one of the three items matched the orientation of the item in the initial display. The duration of the visual task, and the test display of the working memory task, were manipulated across the two types of blocked designs: start-stop and continuous block designs.

**Figure 1.**
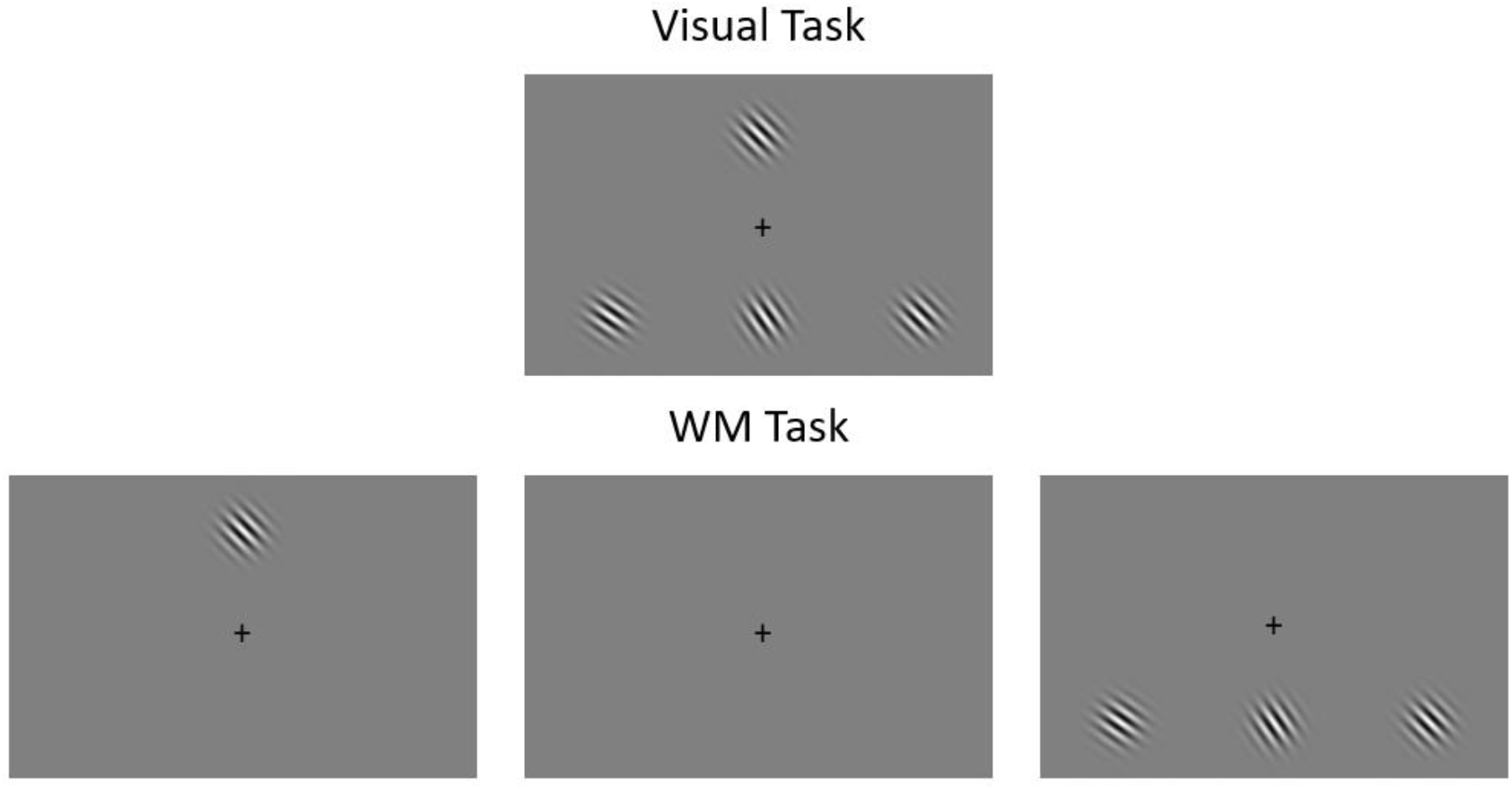
Schematic of the two tasks used in the current study. During fMRI scanning, participants completed a visual task (i.e., perceptual matching task; upper panel) and a working memory task (i.e., delayed match-to-sample; lower panel). The angular difference between the three test items was determined separately for each participant and condition using a staircase procedure. Note that when presented in the start-stop block design, the length of the visual task display was fixed at 4000ms, while the length of the test display in the working memory task was fixed at 3000ms. In the continuous block design, these durations were variable, as the trials ended upon the response of the participant. WM: working memory.

#### Blocked designs

Two types of blocked design were used; both designs involved 36-s task blocks, interleaved with 20-s rest blocks. In *start-stop* blocks, a fixed number of trials of equal length were presented during each block, following the timing parameters of Garrett, Kovacevic, et al. (2011). In this design, a participant’s response in a trial had no bearing on the timing (i.e., after their response, they waited until the next trial). In the visual task, the trial length was 4000ms, and 6 trials were presented in a block (48 trials in total). In the working memory task, the trial length was 7000ms, with 3000ms allotted to responding to the test display. Four trials were presented in a block (32 trials in total). For both visual and working memory tasks, the inter-trial interval was 2000ms.

In *continuous* blocks, all trials terminated upon a participant’s response and, after a brief delay (500ms), proceeded onto the next trial. Because the total block duration was set at 36s, it was possible that a block could end during an experimental trial. In such cases, the trial would terminate prior to a participant’s response and consequently was not included in the calculation of behavioural measures for that condition.

Each task/blocked design combination was presented in a separate functional run, making for four functional task runs. The order of runs was counterbalanced across participants.

#### Staircase procedure

To ensure that all participants performed the visual and working memory tasks at a comparable difficulty level, and to minimise any possible differences in difficulty between tasks involving start-stop and continuous timing parameters, each participant completed a staircase procedure for each task/blocked design combination prior to entering the scanner. The adaptive procedure used was a three-down, one-up algorithm that sought to determine a threshold (angular difference between the orientation of the three test items; see Figure 1) that would result in 80% accuracy for each participant.

#### Behavioural data

For each of the four task/blocked design combinations, three behavioural measures are reported: accuracy (proportion correct), mean response time (meanRT), and response time variability (SD_RT_). Following Garrett, Kovacevic, et al. (2011, 2013), response times were trimmed to exclude response times less than 150ms, and those that were more than three SDs away from the within-subject mean. Missing data (59/6157 trials) were then imputed using the Missing Data Imputation Toolbox (Folch-Fortuny et al., 2016). In addition, SD_RT_ was calculated by first regressing out age, block, trial and all their interactions from raw response times, and then calculating the SD of the residuals separately for each condition (Garrett et al., 2011, 2013).

#### MRI acquisition

MRI data were acquired on a 3-Tesla Siemens MAGNETOM Skyra scanner. A magnetisation-prepared rapidly-acquired gradient echo (MP-RAGE) sequence was used to acquire whole-brain anatomical images. A T2*-weighted echo planar imaging (EPI) sequence was used to acquire whole-brain functional images with 28 interleaved axial slices (5mm thick) acquired in an interleaved fashion (TR = 2000ms, TE = 30ms, FOV = 200mm, flip angle 70°). Four task functional runs were collected, each consisting of 224 volumes. Following the second functional run, a Siemens standard double-echo field map sequence (TR = 577ms; TE = 4.92 and 7.38ms) was used to acquire a set of field maps. A resting-state scan was also collected following the task runs; these data are extraneous to the current study and will not be discussed further.

Task stimuli (black and white Gabor patches displayed on a grey background) were displayed on an MR-compatible BOLD monitor (Cambridge Research Systems, Rochester, U.K.) and viewed with a mirror incorporated into the Siemens 32-channel head coil. Presentation^®^ software (Version 15.0, Neurobehavioral Systems, Inc., Berkeley, CA, www.neurobs.com) was used for the generation and presentation of stimuli. Responses were made using the right three middle fingers on a Pyka MR-compatible fibre optic response box (Current Designs Inc., Philadelphia, P.A.).

#### MRI preprocessing

Preprocessing of MRI data was performed using fMRIPprep 1.1.6 (Esteban et al., 2019), which is based on Nipype 1.1.2 (Gorgolewski et al., 2011, 2018).

#### Anatomical data preprocessing

The T1-weighted (T1w) image was corrected for intensity non-uniformity (INU) using N4BiasFieldCorrection in ANTs 2.2.0 (Tustison et al., 2010), and used as T1w-reference throughout the workflow. The T1w-reference was then skull-stripped using antsBrainExtraction.sh (ANTs 2.2.0), using OASIS as target template. Brain surfaces were reconstructed using recon-all in FreeSurfer 6.0.1 (Dale et al., 1999), and the brain mask estimated previously was refined with a custom variation of the method to reconcile ANTs-derived and FreeSurfer-derived segmentations of the cortical gray-matter of Mindboggle (Klein et al., 2017). Spatial normalisation to the ICBM 152 Nonlinear Asymmetrical template version 2009c (Fonov et al., 2009) was performed through nonlinear registration with antsRegistration (Avants et al., 2008), using brain-extracted versions of both T1w volume and template. Brain tissue segmentation of cerebrospinal fluid (CSF), white-matter (WM) and grey-matter (GM) was performed on the brain-extracted T1w using FAST in FSL 5.0.9 (Zhang et al., 2001).

#### Functional data preprocessing

For each of the four functional (BOLD) runs, the following preprocessing was performed. First, a reference volume (BOLD reference) and its skull-stripped version were generated using a custom methodology of fMRIPrep. A deformation field to correct for susceptibility distortions was estimated based on a field map that was co-registered to the BOLD reference, using a custom workflow of fMRIPrep derived from D. Greve’s epidewarp.fsl script and further improvements of HCP Pipelines (Glasser et al., 2013). Based on the estimated susceptibility distortion, an unwarped BOLD reference was calculated for a more accurate co-registration with the anatomical reference. Head-motion parameters with respect to the BOLD reference (transformation matrices, and six corresponding rotation and translation parameters) were estimated before any spatiotemporal filtering using mcflirt (Jenkinson et al., 2002). BOLD runs were slice-time corrected using 3dTshift from AFNI (Cox & Hyde, 1997). BOLD time-series were resampled onto their original, native space by applying a single, composite transform to correct for head-motion and susceptibility distortions. These resampled BOLD time-series will be referred to as preprocessed BOLD in original space, or just preprocessed BOLD. The BOLD reference was then co-registered to the T1w reference using bbregister (FreeSurfer), which implements boundary-based registration (Greve & Fischl, 2009). Co-registration was configured with nine degrees of freedom to account for distortions remaining in the BOLD reference.

Automatic removal of motion artifacts using ICA-AROMA (Pruim et al., 2015) was performed on the preprocessed BOLD time-series in MNI space after a spatial smoothing with an isotropic, Gaussian kernel of 6mm FWHM (full-width half-maximum). Corresponding non-aggressively denoised runs were produced after such smoothing^1^. Additionally, the aggressive noise-regressors were collected and placed in the corresponding confounds file. Each BOLD time-series was resampled to MNI152NLin2009cAsym standard space, generating a preprocessed BOLD run in MNI152NLin2009cAsym space.

In addition to the denoising performed by ICA-AROMA, we calculated several confounding time-series based on the preprocessed BOLD, which we then regressed out of each voxel’s time-series: framewise displacement (FD), DVARS, three region-wise global signals, and a set of physiological (CompCor) regressors. FD and DVARS were calculated for each functional run, using their implementations in Nipype (following the definitions by Power et al., 2014). The three region-wise global signals extracted were from within the CSF, the WM, and the whole-brain masks. The set of physiological regressors to allow for component-based noise correction were extracted using CompCor (Behzadi et al., 2007). Principal components were estimated after high-pass filtering the preprocessed BOLD time-series (using a discrete cosine filter with 128s cut-off) for the two CompCor variants: temporal (tCompCor) and anatomical (aCompCor). Six tCompCor components were then calculated from the top 5% variable voxels within a mask covering the subcortical regions. This subcortical mask was obtained by heavily eroding the brain mask, which ensures it does not include cortical GM regions. For aCompCor, six components were calculated within the intersection of the aforementioned mask and the union of CSF and WM masks calculated in T1w space, after their projection to the native space of each functional run (using the inverse BOLD-to-T1w transformation). Note that only the first five aCompCor components were used as regressors.

The head-motion estimates calculated in the correction step were also placed within the corresponding confounds file. All resamplings were performed with a single interpolation step by composing all the pertinent transformations (i.e., head-motion transform matrices, susceptibility distortion correction, and co-registrations to anatomical and template spaces). Gridded (volumetric) resamplings were performed using antsApplyTransforms (ANTs), configured with Lanczos interpolation to minimise the smoothing effects of other kernels (Lanczos, 1964). Non-gridded (surface) resamplings were performed using mri_vol2surf (FreeSurfer). Many internal operations of fMRIPrep use Nilearn (Abraham et al., 2014), principally within the BOLD-processing workflow. For more details of the pipeline see fmriprep.readthedocs.io/en/latest/workflows.html.

#### Behavioural data analysis

For each behavioural measure (accuracy, meanRT, and SD_RT_), confirmatory 2 x 2 repeated measures ANOVAs were run in JASP v0.13.01. The two factors were *task* (visual/working memory) and blocked *design* (start-stop/continuous). In addition, planned comparisons were run between start-stop and continuous designs separately for visual and working memory tasks.

#### Calculation of SD_BOLD_ and mean_BOLD_

Raw mean_BOLD_ was calculated for each voxel using the PLSgui package (https://www.rotman-baycrest.on.ca/index.php?section=84). First, the mean percent signal change (relative to the onset of the block) for each block, and then these block-means are averaged across all blocks (McIntosh et al., 2004; McIntosh & Lobaugh, 2004). Absolute mean_BOLD_ was also derived for each voxel by converting raw mean_BOLD_ to absolute values, thus reflecting the mean percent change from zero irrespective of whether the changes from the block onset were positive (activation) or negative (deactivation) in nature.

SD_BOLD_ values were calculated for each voxel using the Variability Toolbox (https://github.com/LNDG/vartbx), which adopts the procedure outlined in a number of papers investigating BOLD variability (e.g., Garrett et al., 2014; Garrett, Kovacevic, et al., 2011, 2013; Grady & Garrett, 2014, 2018). To account for scanner drift across the run, mean signal within each block was normalised to 100. Next, the block mean was subtracted from signal intensity at each time-point within a block. Finally, all task blocks within a run were concatenated, and the SD was calculated on this concatenated time-series (see Figure 2).

**Figure 2.**
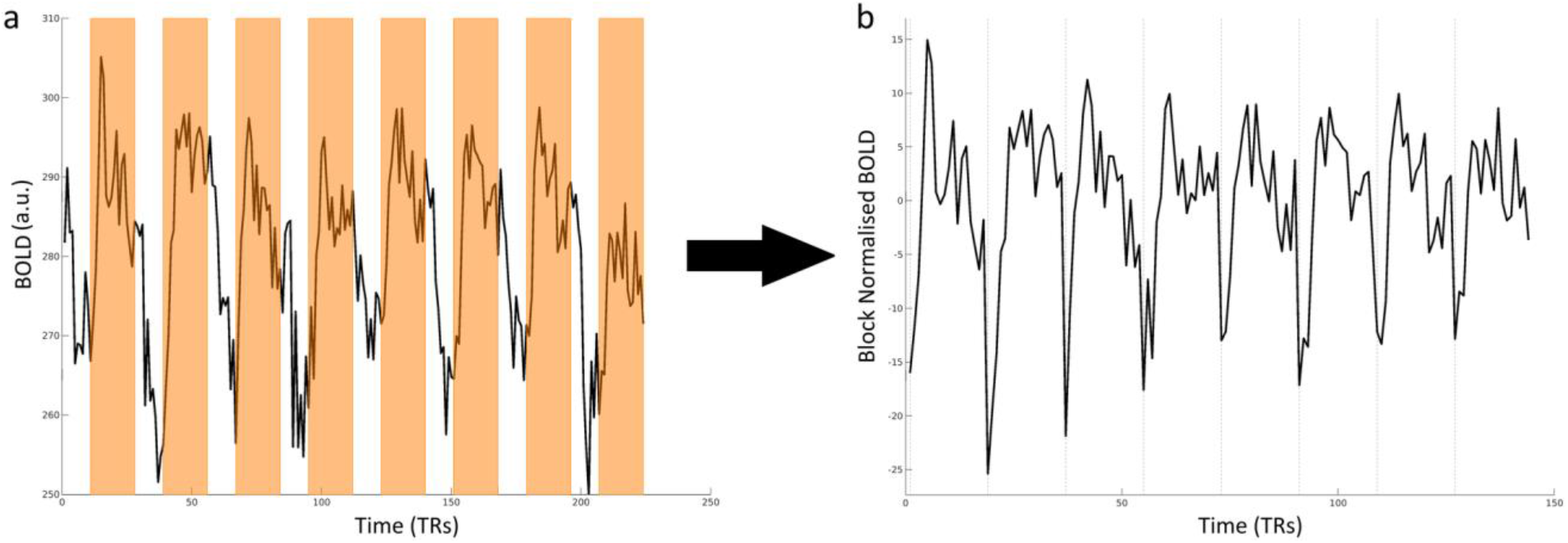
An illustration of how task-related BOLD data were prepared for SD_BOLD_ calculations using the variability toolbox in SPM. The left panel (a) shows a voxel time-series (34 −70 −20) from a single participant for the visual task/continuous blocked design run; periods corresponding to 36-s task blocks are shaded orange; the unshaded blocks were 20-s rest blocks. The right panel (b) shows the time-series produced by the variability toolbox (after block normalisation), on which standard deviations were calculated.

#### Partial least-squares analyses

Many hypotheses in the current study were tested using partial least squares (PLS), a technique commonly used in BOLD variability studies (Boylan et al., 2020; Garrett et al., 2014; Garrett, Kovacevic, et al., 2011, 2013; Roberts et al., 2020). PLS is a multivariate technique that examines pattern of covariances across brain and design (i.e., conditions/behaviours) matrices, resulting in latent variables that comprise sets of voxels associated with contrasts of tasks and/or behaviours (Krishnan et al., 2011; McIntosh et al., 2004; McIntosh & Lobaugh, 2004). Importantly, because all voxels, tasks, and behaviours are included in a single multivariate analysis, no need for multiple comparisons is required.

Each PLS analysis used the following general procedure. First, as described above we calculated BOLD data (mean or SDs) during experimental task blocks for all voxels within a grey matter mask. The mask was created by thresholding FSL’s MNI grey matter tissue probability map (threshold = 0.4), and binarising the thresholded image. The BOLD data were then flattened (i.e., converted into a vector), and all participants’ data were then “stacked” into a group brain matrix. The brain matrix was then cross-correlated with a matrix containing information about the contrast of tasks or behaviours (dependent on the analysis), producing a cross-block matrix that was submitted to singular value decomposition, producing latent variables that maximally accounted for the relationship between the brain and task/behaviour matrices. While PLS can be run in an exploratory fashion, in which latent variables are defined by contrasts that maximally account for the relationship between brain and tasks/behaviours, all PLS analyses reported here are “non-rotated”, in which an hypothesis-driven contrast was coded into the design.

The latent variables produced by PLS are comprised of: i) a singular value indicating the amount of covariance explained by the latent variable; ii) a linear contrast expressing differences and commonalities between tasks/behaviours and iii) a vector of voxel weights (or “salience”) expressing the magnitude and direction of each voxel’s relationship to the contrast of tasks/behaviours. Brain scores, a weighted average of all voxel saliences, were calculated for each task/behaviour for each participant, and provide a metric of how strongly the brain pattern of a given latent variable was expressed by that participant.

The statistical significance of each latent variable was assessed by permutation testing (1000 permutations). For task PLS (testing differences between tasks), on each permutation, all participants’ data were reassigned to tasks, the analysis was rerun, and a new singular value obtained. Significance indicated the probability of the number of times the singular value from the permuted data exceeded the actual singular value obtained (McIntosh et al., 1996); a threshold of *p* < .05 was used. For behaviour PLS (testing differences/commonalities in brain-behaviour correlations), participants’ behavioural data were shuffled on each permutation.

Bootstrapping estimation of the standard error (SE) was used to determine the reliability of voxel saliences (1000 bootstraps). Voxels with bootstrap ratios (i.e., the ratio of each voxel’s salience to its standard error; BSR) greater than ±2.58 (equivalent to *p* <.01) were considered to represent reliable voxels.

#### The relationship between SD_BOLD_ and mean_BOLD_

To determine the relationship between SD_BOLD_ and mean_BOLD_, a confirmatory analysis was performed assessing the within-subject, voxelwise correlation between i) SD_BOLD_ and *raw* mean_BOLD_ and ii) SD_BOLD_ and *absolute* mean_BOLD_. Voxelwise Pearson correlation coefficients were calculated for each task/blocked design combination between SD_BOLD_ and *raw* mean_BOLD_, and between SD_BOLD_ and *absolute* mean_BOLD_, yielding eight correlations per participant. Correlation coefficients were then Fisher z-transformed. For each of the eight z-transformed correlations, bootstrapped confidence intervals were calculated; intervals that included zero were taken to indicate no relationship between SD_BOLD_ and mean_BOLD_. This estimation approach was supplemented by a series of voxelwise one-sample *t*-tests to test whether z-transformed correlations were significantly different from zero. Finally, a series of paired-sample *t*-tests were performed to determine whether, for any of the task/design combinations, the voxelwise correlation of SD_BOLD_ with *raw* mean_BOLD_ differed from that with *absolute* mean_BOLD_. All *t*-tests were Bonferroni corrected for multiple comparisons. All t-tests were computed in JASP v0.13.01.

To further interrogate the robustness of the relationship between SD_BOLD_ and mean_BOLD_, we performed exploratory analyses that assessed *across-subject* correlations at each voxel. That is, for each *voxel*, we ran across-subject correlations between SD_BOLD_ and raw mean_BOLD_, and between SD_BOLD_ and absolute mean_BOLD_, for each task/design combination. We generated histograms of correlation coefficients to explore whether there was a systematic shift to the right in correlation coefficients (i.e., more positive correlations) when SD_BOLD_ was correlated with absolute mean_BOLD_, relative to when SDBOLD was correlated with raw mean_BOLD_ values. In addition, we generated spatial maps to illustrate the spatial extent of voxels showing significant correlations of SD_BOLD_ with both raw and absolute mean_BOLD_.

#### SD_BOLD_ differences between start-stop and continuous designs

A confirmatory non-rotated task PLS analysis was performed to test for systematic differences in SD_BOLD_ between the two forms of blocked design used; we predicted that start-stop design should result in increased BOLD variability relative to continuous designs. In addition, a series of exploratory analyses were performed to investigate the effect of mean_BOLD_ on these effects.

#### Relationship between SD_BOLD_ and response times

A confirmatory, non-rotated behaviour PLS analysis was run to directly compare the brain-behaviour correlations between the start-stop and continuous block designs. That is, meanRT and SD_RT_ were used as behaviours, and we directly contrasted the magnitude of brain-behaviour correlations for start-stop designs against those for continuous designs. This analysis was complemented by exploratory analyses in which we assessed the magnitude of brain-behaviour correlations separately for each type of block design.

#### Modelling BOLD time-series using response time patterns

Finally, we ran an exploratory analysis testing whether simulated BOLD time-series incorporating the specific pattern of response times within a run is predictive of participants’ actual BOLD time-series. If toggling between on- and off-task states in start-stop designs contributes to variability of the BOLD signal, we hypothesised that taking into account each participant’s specific pattern of response times during a run should aid in predicting the time-series of voxels in start-stop designs. To test this, we used the R-package NeuRosim (Welvaert et al., 2011) to create simulated time-series for all trials across the visual and working memory task start-stop runs. Specifically, we specified the onset and response time of each trial, using this time between the two as the trial duration, which was convolved with a canonical hemodynamic response function (HRF). Task blocks were then concatenated to form a simulated time-series (see Figure 3a). This *response-time regressor* was used as a predictor for the actual concatenated time-series of experimental blocks (i.e., the time-series on which SD_BOLD_ was calculated) in a voxelwise general linear model; the resulting voxel beta-coefficients indicated how well response times as trial durations predicted each voxel’s actual time-series, and were used for subsequent analyses. In addition, we repeated the same process for three other simulated time-series, shown in Figure 3b-d: a *whole-block regressor* in which onsets were specified for each block, and the entire block length was used as duration; a *whole-trial regressor*, in which trial onsets were specified and the trial-duration was specified as the duration irrespective of when a response was made; and a *stick-function regressor*, in which the trial onsets were used, and the trial-duration was specified as 0.

**Figure 3.**
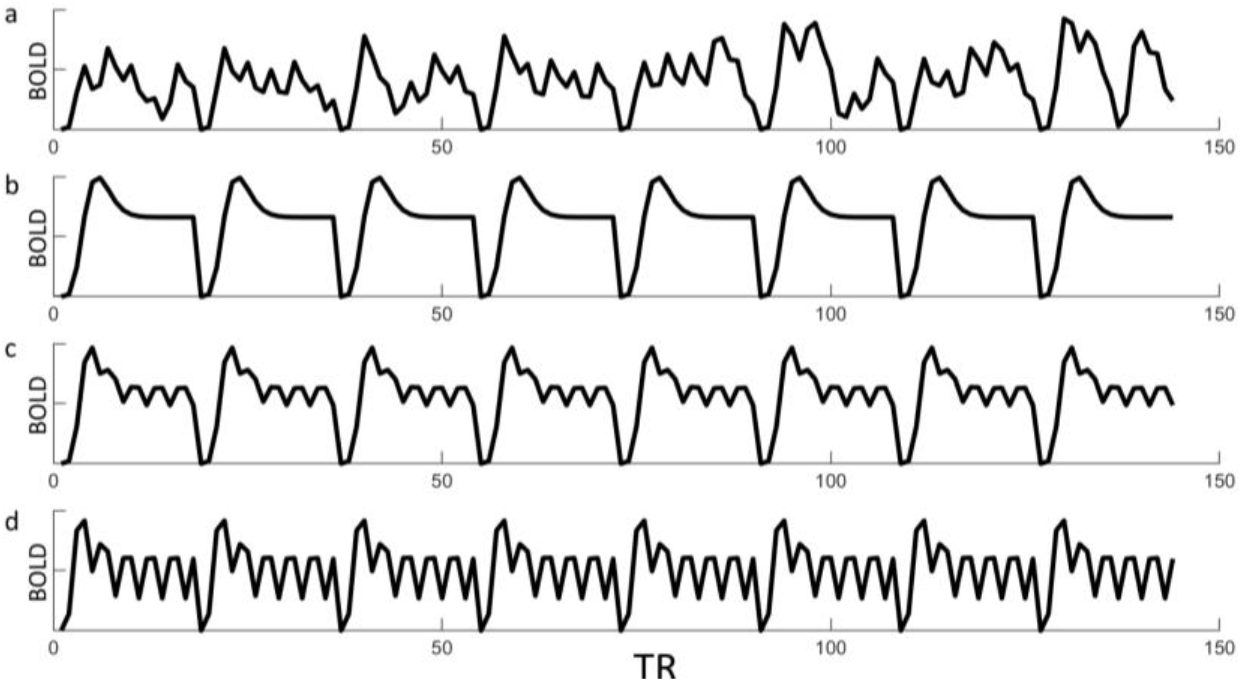
Simulated time-series regressors for the start-stop design generated using a: trial onsets as onsets and response times as durations; b: block onsets as onsets and block lengths as durations; c: trial onsets as onsets and whole trials as durations; d: trial onsets as onsets and durations of 0 (i.e., stick functions at task onsets).

Non-rotated task-PLS analyses were then run on the beta-coefficients, separately for visual and working memory tasks, contrasting the predictive strength of the response-time regressors to all other regressors (whole-block, whole-trial, and stick-function time-series). We predicted that the response-time regressor should be a better predictor of the actual data and thus, relative to the other regressors, it should produce increased effects in task-positive regions (e.g., visual, motor, dorsal attention networks) and decreased effects in the default-mode network due to this network showing deactivations in both tasks (i.e., the response-time regressor should produce more positive and more negative beta-coefficients in task-positive and default-mode regions, respectively).

## Results

### Behavioural Results

A summary of behavioural results (accuracy, meanRT and SD_RT_) are displayed in Figure 4. No significant main effects or interactions were found for accuracy (all *p*’s > 0.47). For meanRT, there was a main effect of *task*, F(1,22) = 65.73,*p* < .001, with responses in the working memory task being faster than in the visual task (mean difference = 413ms, 95% CI [308, 519]). The main effect of *design* was also significant, F(1,22) = 49.02, *p* < .001, with responses in the continuous blocks being faster than start-stop blocks (mean difference = 298ms, 95% CI [210, 387]). Bonferroni-corrected planned comparisons showed that this design effect was apparent for response times in both the visual (*p* <.001; mean difference = 448ms, 95% CI [346, 550]) and working memory tasks (*p* = .004; mean difference = 148ms, 95% CI [59, 237]). Finally, the *task* x *design* interaction was significant, *F*(1, 22) = 71.93, *p* < .001, and was driven by the difference between start-stop and continuous designs being more pronounced in the visual than in the working memory task (see Figure 4b).

**Figure 4.**
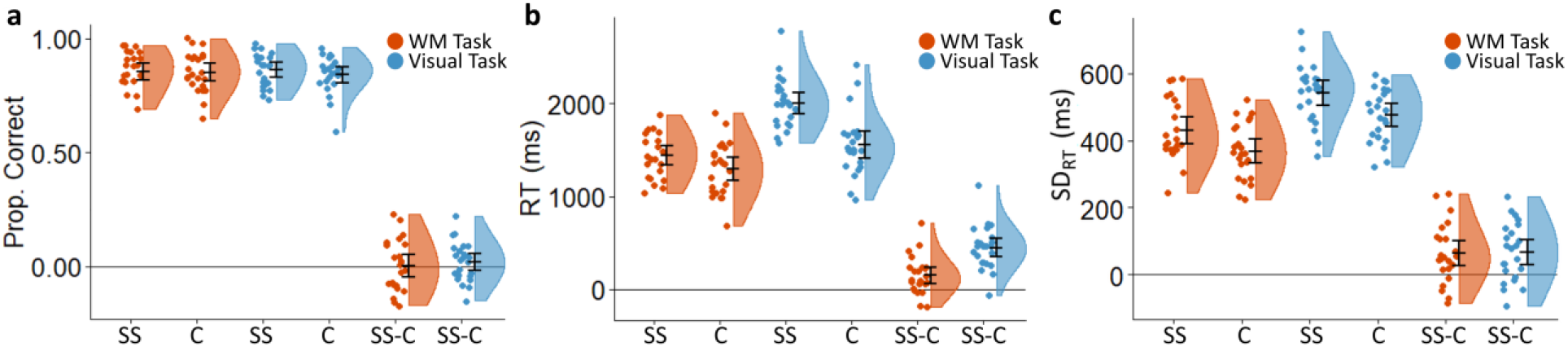
Summary statistics for a) accuracy, b) mean response time, and c) response time variability. Error bars indicate 95% confidence intervals. C: continuous; prop: proportion; RT: response time; SS: start-stop; SS-C: difference scores between start-stop and continuous designs; WM: working memory.

A similar pattern of results were found for SD_RT_. The main effect of *task* was significant, *F*(1, 22) = 41.21, *p* < .001, with response times being more variable in the visual task relative to the working memory task (mean difference = 110ms, 95% CI [74, 146]). The significant main effect of *design, F*(1, 22) = 20.90, *p* < .001, was driven by more variable responses in the start-stop design (mean difference = 65ms, 95% CI [35, 94]). Bonferroni-corrected planned comparisons showed that the effect of *design* was significant for both visual (*p* = .002; mean difference = 67ms, 95% CI [29, 104]) and working memory tasks (*p* = .006; mean difference = 63ms, 95% CI [25, 102]). However, unlike meanRT, there was no significant task x block design interaction (*p* = .88).

### The relationship between SD_BOLD_ and mean_BOLD_

A summary of voxelwise correlations are displayed in Figure 5. Bonferroni-corrected one-sample *t*-tests for all four task/design combinations showed no evidence for a reliable voxelwise relationship between SD_BOLD_ and raw mean_BOLD_: all *p*’s > .07. There was, however, strong evidence for a linear relationship between SD_BOLD_ and absolute mean_BOLD_ across all conditions: all *p*’s < 6.10E-18. Finally, paired-sample *t*-tests showed significant differences between the two sets of voxelwise correlations for all task/design combinations, with absolute mean_BOLD_ correlations being greater than raw mean_BOLD_ correlations: all *p*’s < 4.49E-09. Together, these confirmatory analyses provide strong evidence that the relationship between means and SD is one in which SD is strongly associated with *absolute* mean signal change, but not reliably so with raw mean_BOLD_.

**Figure 5.**
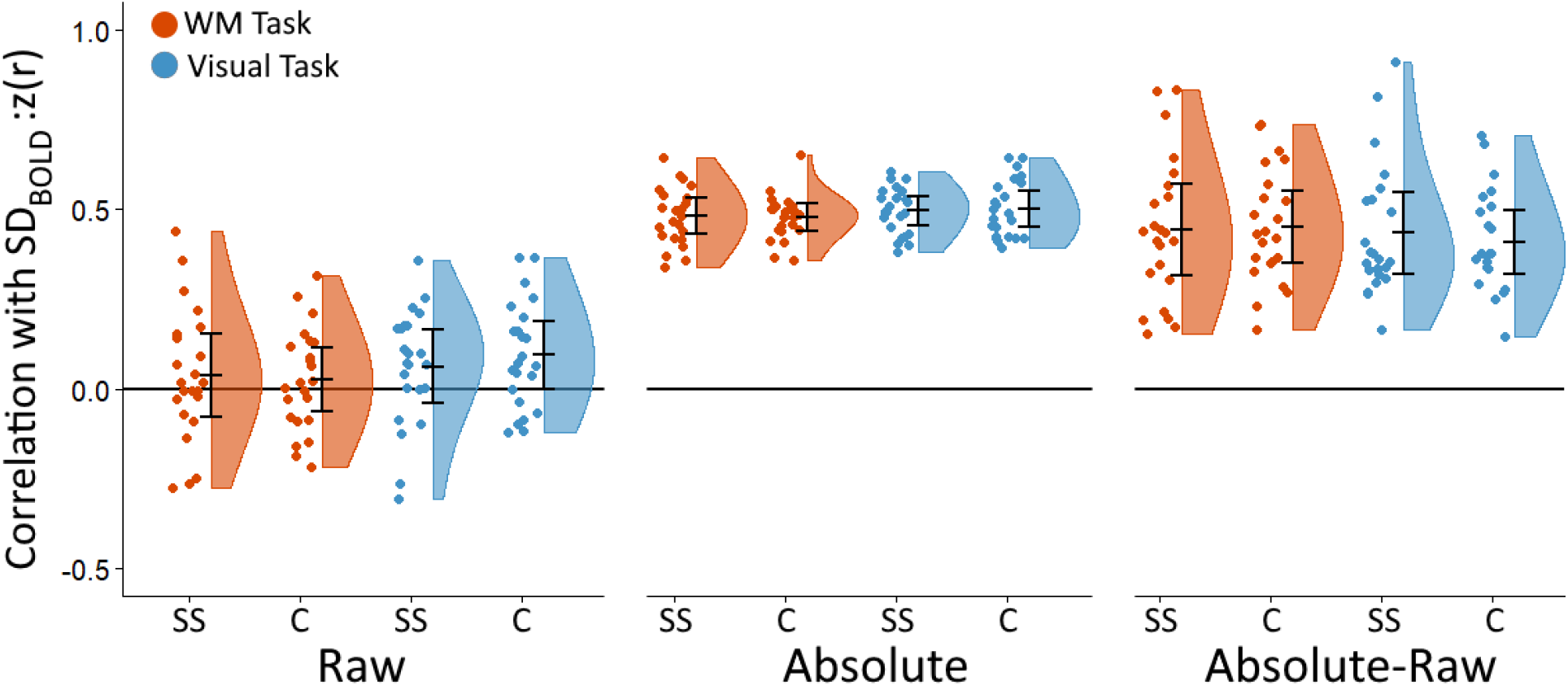
Z-scored voxelwise correlations showing voxelwise relationship for all conditions between SD_BOLD_ and raw mean_BOLD_ (left set of plots) and between SD_BOLD_ and absolute mean_BOLD_ (middle set of plots). Also shown are the difference scores for the two sets of correlations (right set of plots). Error bars indicate 95% confidence intervals, corrected for multiple comparisons. C: continuous design; SS: start-stop design; WM: working memory.

In addition, as shown in Figure 6, the distribution of across-subject correlation coefficients showed a systematic increase in correlation coefficients for the relationship between SD_BOLD_ and absolute mean_BOLD_. This effect was reflected in a substantial increase in the number of voxels showing a significant (i.e., *p* < .05) positive relationship between SD_BOLD_ and absolute mean_BOLD_.

**Figure 6.**
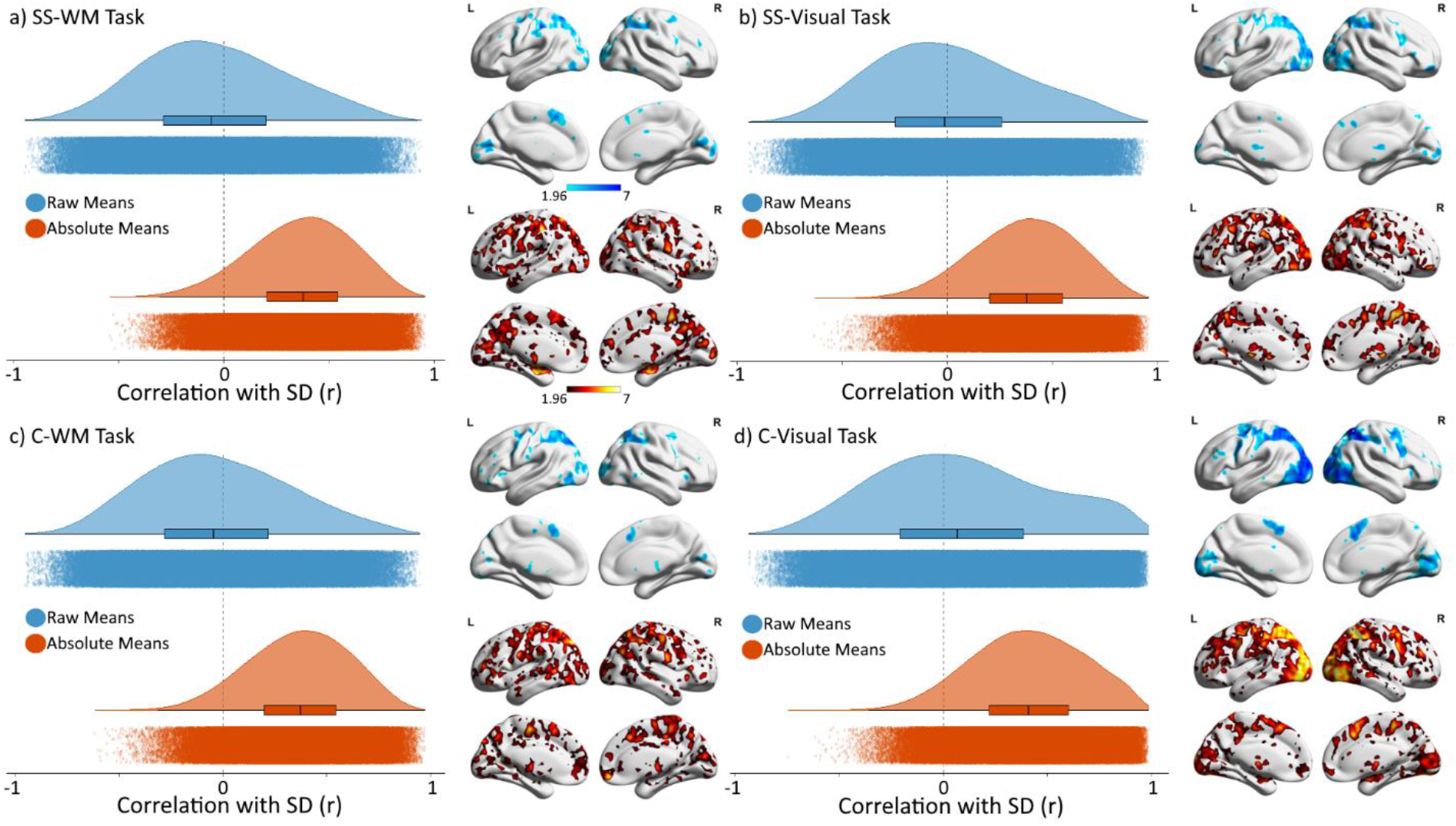
Histograms showing the distribution of across-subject correlation coefficients across all voxels for the four task/design combinations. Also shown are spatial maps of voxels showing a significant positive association for the two correlation types, thresholded at *z* > = 1.96 (*p* < .05). Maps are uncorrected for multiple comparisons, and are shown for descriptive purposes. Blue histograms and clusters denote the relationship between SDBOLD and raw mean_BOLD_; orange histograms and clusters denote the relationship between SDBOLD and absolute mean_BOLD_. C: continuous design; SS: start-stop design; WM: working memory.

### SD_BOLD_ differences between start-stop and continuous designs

We predicted that start-stop block designs would induce an increase in BOLD variability, relative to continuous designs, due to the potential toggling between on- and off-task states during start-stop blocks. A confirmatory, non-rotated task-PLS that directly contrasted the two block designs failed to show any evidence for the hypothesised effect (*p* = .20).

We reasoned that this lack of difference in SD_BOLD_ between the start-stop and continuous designs may have, in fact, been obscured by differences in mean_BOLD_ between the designs. Specifically, if the continuous designs resulted in greater (absolute) mean signal change than the start-stop designs, and if absolute mean changes show reliable associations with BOLD variability as we have demonstrated in previous analyses, differences in mean BOLD signal could feasibly mask any differences in BOLD variability between the two designs related to whether the design induces toggling between task states. To test this hypothesis, we initially ran an exploratory non-rotated task PLS analysis, which showed reliable whole-brain differences in mean BOLD signal between the designs (*p* = .026), with task-positive regions (visual, dorsal attention and frontoparietal control networks) showing greater mean activation and default-mode network regions (e.g., precuneus) showing greater mean deactivations in the continuous designs.

Given the reliable differences in mean_BOLD_ between start-stop and continuous designs, and the clear linear relationship between SD_BOLD_ and absolute mean_BOLD_ described earlier, we again tested for the predicted differences in SD_BOLD_ between start-stop and continuous designs after controlling for differences in mean BOLD signal between the two task designs. We did this in two ways. We first used a *partial block method*, in which the first three TRs of each block were simply removed from the concatenated, block-normalised time-series, and SD was recalculated. This was done given evidence to suggest that these initial TRs can be problematic in the standard block-normalised time-series from which SD is typically derived; as can be seen in Figure 2b, a voxel that reliably activates (or deactivates) to task blocks exhibits sharp increases (or decreases) in BOLD signal at block onsets that contribute to *both* the mean and SD for that voxel. Thus, we reasoned that removing these volumes would remove the confounding effects of mean_BOLD_, while still leaving a long enough time-series to capture the moment-to-moment variability associated with each condition. Our second approach was a *task regression method*, in which the task design (box-car function) was regressed out from each voxel’s time-series. This method removes the increases (or decreases) in signal associated with block onsets, while maintaining signal variability subsequent to the initial increase (or decrease) in each block. Block normalisation and concatenation of task blocks were then performed, producing time-series on which SD_BOLD_ was calculated.

Non-rotated task PLS on these SD_BOLD_ data showed evidence for start-stop tasks resulting in increased BOLD variability relative to continuous blocked designs for both the partial block (*p* = .010) and task regression (*p* = .041) methods. As shown in Figure 8, the latent variables for both analyses were dominated by voxels from a widespread set of regions, showing the predicted increase in variability in the start-stop designs relative to continuous versions of the task.

**Figure 7.**
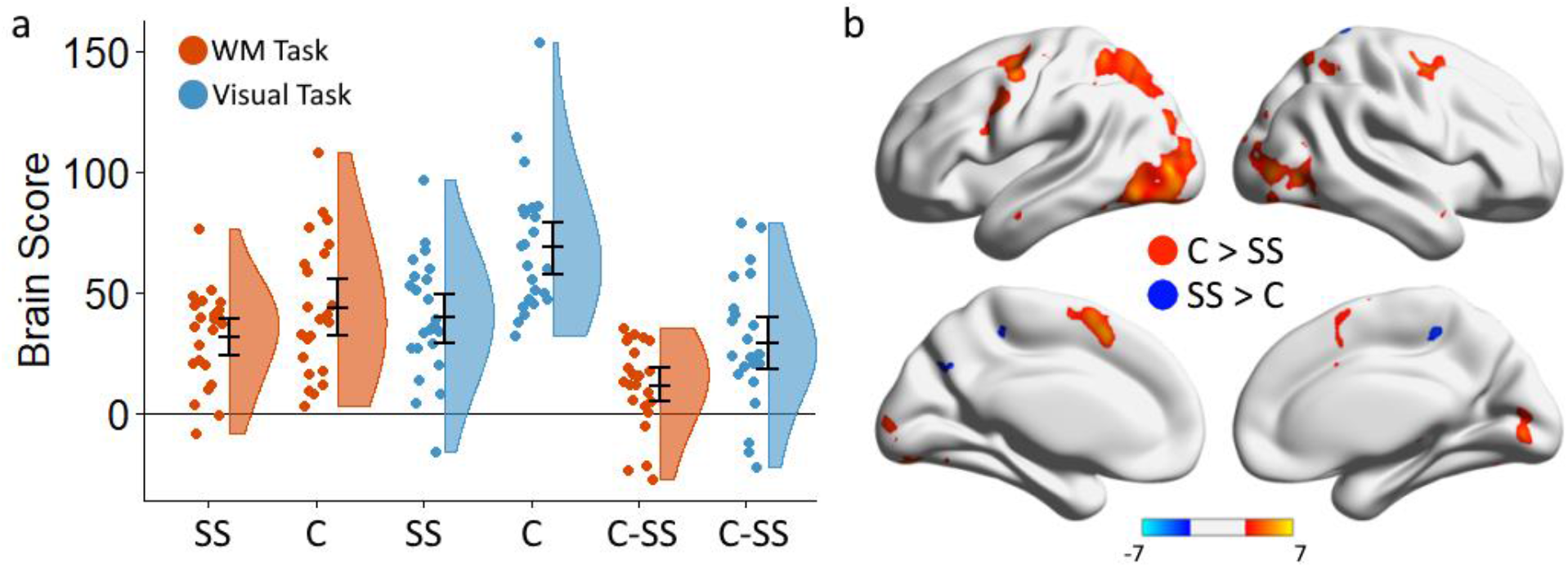
Summary of task PLS showing reliable differences in mean_BOLD_ between continuous and start-stop designs. 7a shows brain score plots for each condition, as well as difference scores between continuous and start-stop conditions for memory and visual tasks. Error bars are bootstrapped 95% confidence intervals of the mean; 7b shows a spatial map of voxels reliably contributing to the latent variable, thresholded at a bootstrap ratio of 2.58 (*p* < .01). C: continuous design; SS: start-stop design.

**Figure 8.**
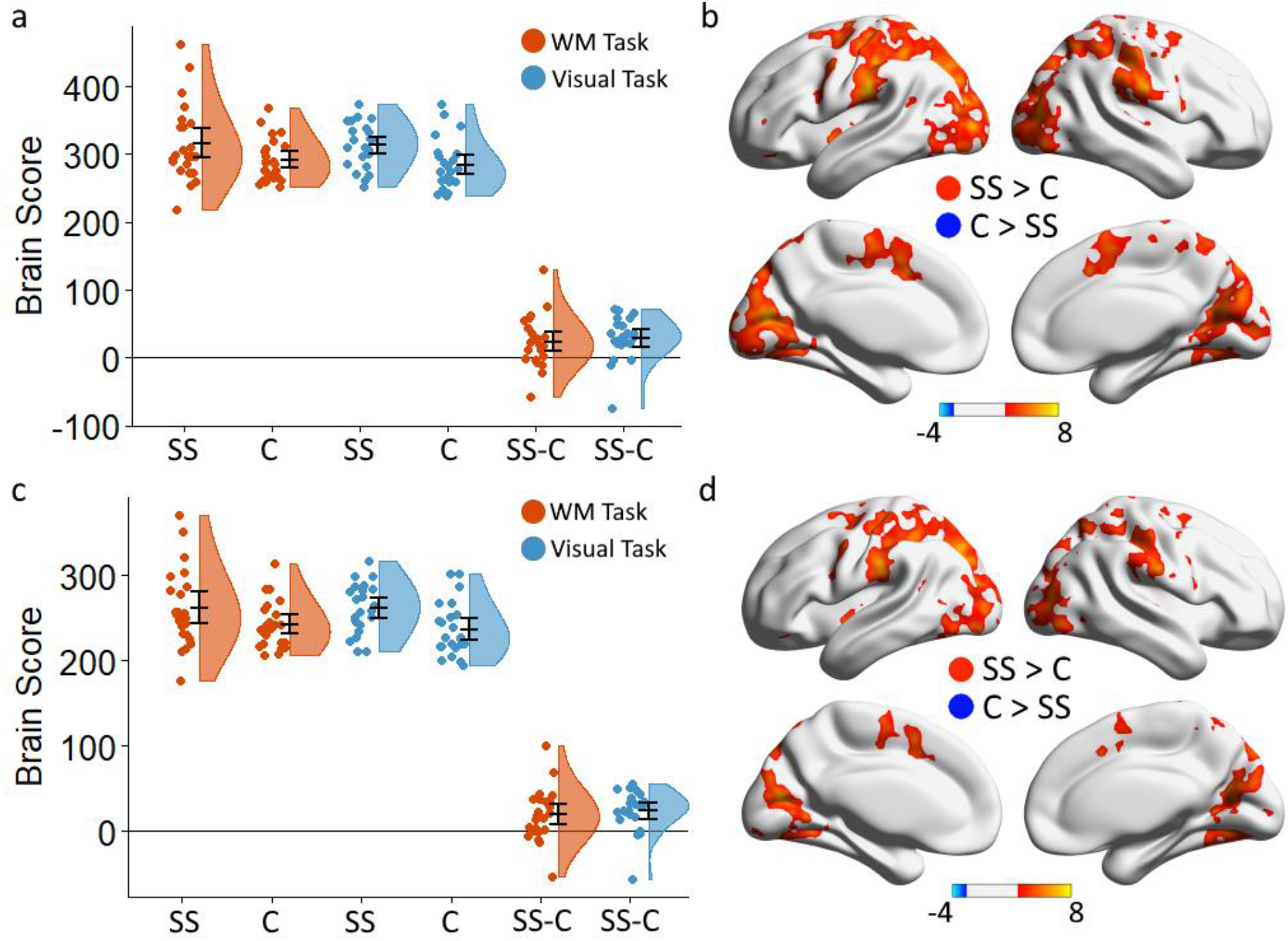
Brain score plots (left) and spatial maps of voxels reliably contributing to latent variable (right), thresholded at a bootstrap ratio of 2.58 (*p* < .01) for non-rotated task PLS analyses contrasting SD_BOLD_ in start-stop and continuous designs. The top panel (a-b) shows the result from SD data computed using the partial block method; the bottom panel (c-d) shows results from SD data computed using the task regression method. Error bars are bootstrapped 95% confidence intervals of the mean. C: continuous; SS: start-stop.

### Relationship between SD_BOLD_ and response times

We predicted that the relationship between response time metrics (meanRT and SD_RT_) and BOLD variability would be greater in the start-stop versions of the task relative to the continuous designs. To this end, we ran a confirmatory, non-rotated behaviour PLS analysis directly contrasting the magnitude of brain-behaviour correlations (i.e., correlations between SD_BOLD_ and RT measures) between the two design types. Permutation tests showed no evidence for this effect (*p* = .310). Exploratory analyses showed that this null effect was also apparent for SD values calculated using the partial block and task-regressed methods described earlier (partial block: *p* = .649; task-regressed: *p* = .571).

To explore the relationship between response time metrics (both meanRT and SD_RT_) and BOLD variability further, we ran non-rotated behavior PLS analyses separately for start-stop and continuous designs to determine if the relationship between response time metrics and SD_BOLD_ was apparent for start-stop designs but not for continuous designs. A significant latent variable (*p* = .041) was produced only for the start-stop designs, showing negative relationships between SD_BOLD_ and both response time metrics in a number of regions across the brain. As shown in Figure 9, however, the correlation profile shows that the relationship between SD_BOLD_ and SD_RT_ for the working memory task did not reliably contribute to the latent variable, as the bootstrapped 95% confidence interval includes zero. Similar results were found for the same analysis run on partial block and task-regressed SD_BOLD_ data (partial block: *p* = .048; task-regressed: *p* = .043). This effect was not, however, apparent for the continuous designs using standard (*p* = .268), partial block (*p* = .266) or task-regressed (*p* = .329) SD_BOLD_ data. Together, these results show some evidence that, in spite of the null result for the confirmatory behaviour PLS analyses that contrasted brain-behaviour relationships between the two block designs, a negative relationship between response time metrics and SD_BOLD_ is present for standard (i.e., start-stop) block designs while no such relationship was found for continuous designs.

**Figure 9.**
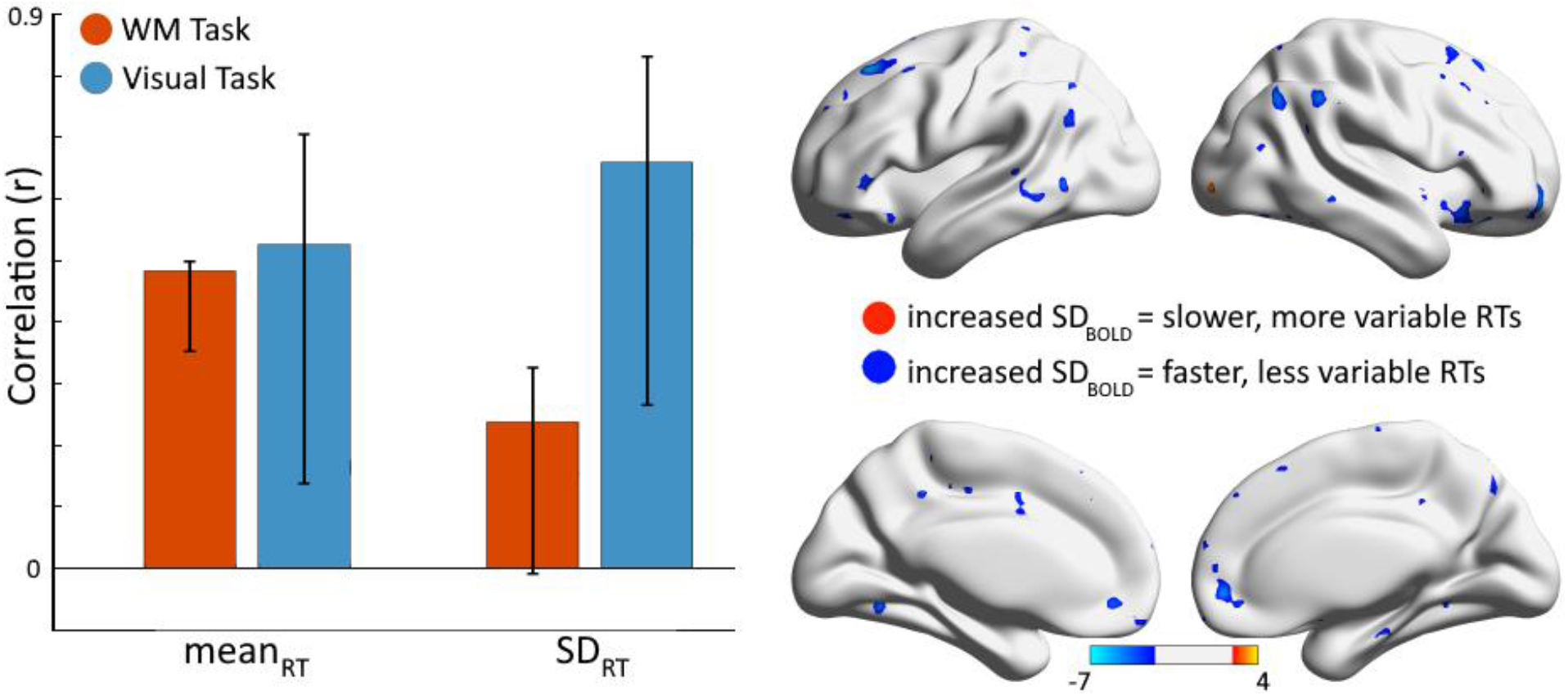
Correlation profile of behaviour PLS analysis showing the relationship between BOLD variability and response times for the start-stop design (left) and the spatial map showing voxels contributing to the latent variable (right), thresholded at a bootstrap ratio of 2.58 (*p* < .01). Warm clusters indicate voxels in which increased SD_BOLD_ is associated with slower, more variable response times (i.e., increased SDRT); cool clusters indicate voxels in which increased SD_BOLD_ is associated with faster, less variable response times (i.e., decreased SD_RT_).

### Using response time patterns to predict BOLD time-series

Finally, we tested if participant-specific response time patterns across functional runs were predictive of the BOLD signal during experimental blocks. Using the simulated BOLD time-series regressors (shown in Figure 3) we predicted that if toggling between on- and off-task states in start-stop designs contributes to variability of the BOLD signal, the response-time regressor would be a better predictor of the actual BOLD data than the other regressors. A series of non-rotated task PLS analyses showed significant differences between the ability of simulated regressors (see Figure 3) to predict the fluctuations of the BOLD signal during start-stop task blocks. Specifically, contrasting the beta-maps of the response-time regressors with maps of whole-block, whole-trial, and stick function regressors produced widespread differences for all contrasts, for both memory and visual conditions (all permutation test *p*’s < .005). Furthermore, as shown in Figure 10a-b, when contrasted with a simple block regressor, the response-time regressor produced higher beta-values in task-positive regions where beta-coefficients were positive (e.g., visual, motor, ventral and dorsal attention networks) and generally lower beta-values for the response-time regressor in default mode regions where beta-coefficients were negative, suggesting that BOLD signal fluctuations during experimental blocks are related to the dynamic pattern of response times of each participant. Similar spatial patterns were found when comparing the response-time regressor to trial duration regressors for the visual condition (Figure 10c), and when comparing the response-time regressor to stick-function regressors for the working memory condition (Figure 10f). However, exceptions to this general pattern occurred in the working memory condition when comparing the response-time regressor to the trial regressor (Figure 10d), and in the visual condition when comparing the response-time regressor to the stick-function regressor (Figure 10e), suggesting that the degree to which a region tracks closely with toggling between on- and off-task states during a block is not uniform across brain regions, and is contingent on the specific timing parameters of experimental trials.

**Figure 10.**
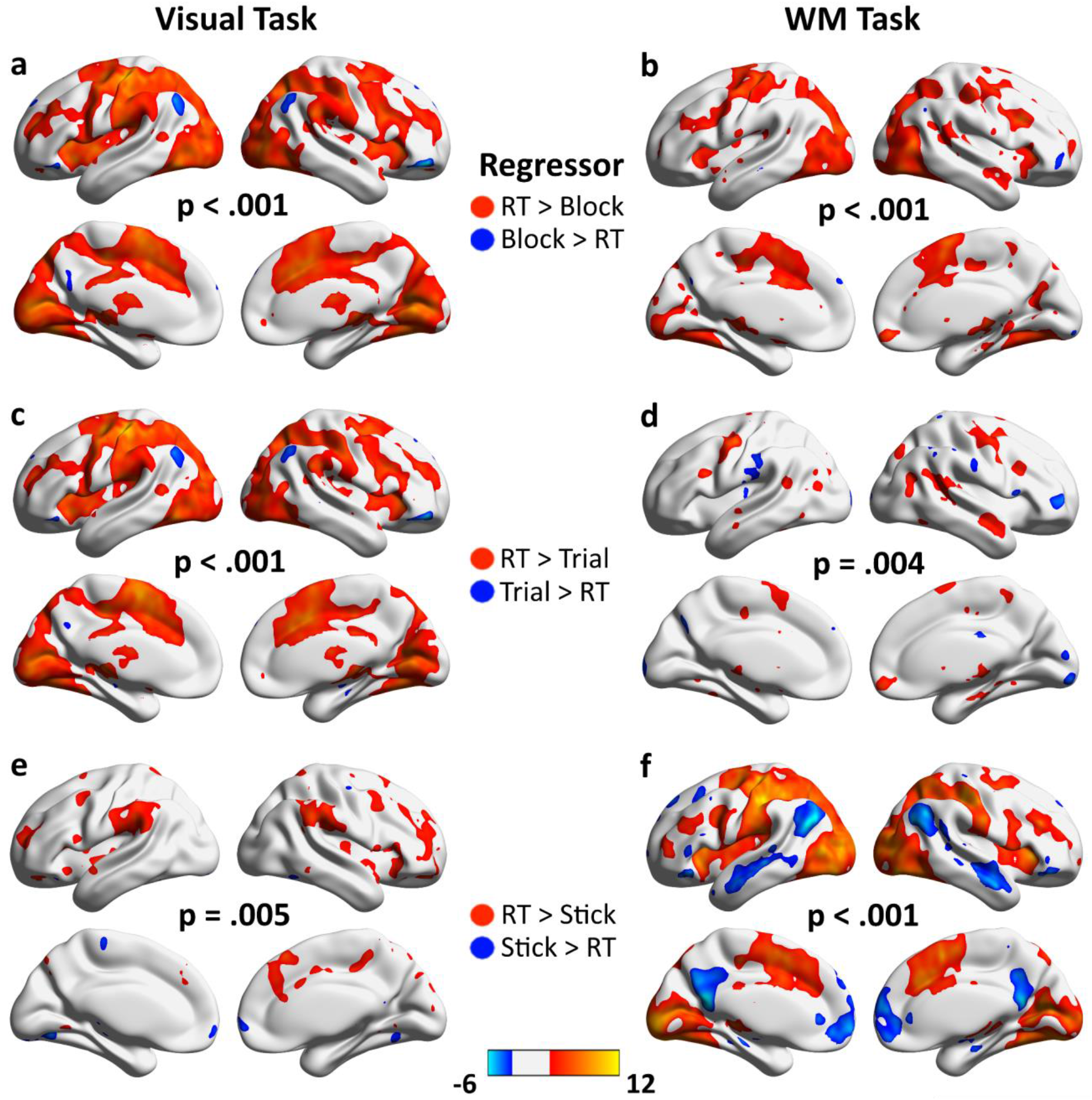
Spatial maps showing voxels in which the response-time regressor produced reliably different beta-coefficients relative to three alternative regressors: a whole-block regressor (a-b), a trial regressor (c-d), and a stick function regressor (e-f). All maps are thresholded at a bootstrap ratio of 2.58 (*p* < .01). *p*-values indicate the result of the PLS permutation test for each analysis. RT: response time; WM: working memory

## Discussion

We investigated two claims commonly made regarding BOLD variability: that SD_BOLD_ is a measure of the BOLD signal that is independent of mean_BOLD_, and that increased SD_BOLD_ captures a property of neural systems that facilitates fast, stable responses during cognitive tasks. The results of the current study shed doubt on both claims. First, correlational analyses across two levels (within- and across-subject) provided evidence that SD_BOLD_ shows a robust linear relationship with absolute mean_BOLD_, suggesting that increased SD_BOLD_ is associated with greater levels of activation or deactivation of the BOLD signal during cognitive tasks. Second, using two forms of blocked design (start-stop and continuous), we provided evidence that many of the SD_BOLD_ effects reported in the literature may have arisen due to the toggling between on- and off-task states that occurs in standard (i.e., start-stop) block designs. In support of this claim, we found that—after controlling for differences in mean_BOLD_—stop-start designs resulted in increased BOLD variability compared to continuous designs in both visual and memory tasks, indicating that switching between task states is a reliable contributor to SD_BOLD_ effects in standard block designs. Next, brain-behaviour correlations showed that increased SD_BOLD_ was associated with faster, less variable responses for start-stop designs. Critically, this effect was not apparent for continuous designs. Finally, we showed that incorporating each participant’s individual response-time patterns during experimental blocks generally led to better prediction of the moment-to-moment fluctuations of the BOLD signal than simply using trial or block onsets as predictors.

Together, these results have broad implications for how to best interpret the findings of studies investigating BOLD variability and its relationship to cognition. First, in task-related fMRI studies, it is not the case that SD_BOLD_ provides a metric of brain activity untethered from traditional, mean-based measures of the BOLD signal. Rather, both measures of the BOLD signal are influenced by sharp increases or decreases at the onset of experimental blocks. It is important to note that the observed relationship between SD_BOLD_ and mean_BOLD_ is likely compounded by the concatenation of experimental blocks in the process of calculating SD_BOLD_. This is because, in voxels that are responsive to the task (i.e., voxels that activate or deactivate), concatenation of experimental blocks produce a steep decline in BOLD signal at the boundary between the end of one block and the beginning of the next (see Figure 2). The process of block normalisation, which occurs prior to block concatenation, has been proposed to mitigate the association between mean and variability effects. Importantly, however, it is not the case that normalising the mean BOLD signal in all blocks to an arbitrary value (e.g., 100) results in all blocks exhibiting the same mean percent signal *change* relative to block onsets, and it is this latter metric to which mean_BOLD_ refers, and which the current results show is associated with SD_BOLD_. The relationship between SD_BOLD_ and mean_BOLD_ described here can, at least in part, account for some aspects of SD_BOLD_ effects previously described in the literature. For example, Garrett and colleagues (2013) reported that a range of externally-driven tasks result in increased variability in both the default-mode and task-positive networks relative to fixation. This stands in stark contrast to the generally reported pattern of DMN mean_BOLD_ deactivation and task-positive mean_BOLD_ activation for these tasks (Smith et al., 2009). However, given the positive relationship between SD_BOLD_ and absolute mean_BOLD_ reported here, it is likely that both mean_BOLD_ deactivations in the DMN and mean_BOLD_ activations in task-positive networks contribute to increased SD_BOLD_ effects across both networks.

While previous research has framed the relationship between cognitive performance—as indexed by response time measures—and SD_BOLD_ as one in which SD_BOLD_ captures a feature of neural systems that result in increased cognitive performance (Garrett, Kovacevic, et al., 2013; Garrett, Samanez-Larkin, et al., 2013), the current results indicate a more parsimonious account that reverses the causal path of this standard interpretation: fast response times induce a more variable BOLD signal during experimental blocks due to toggling between on- and off-task states. This account is bolstered by exploratory analyses showing that individual response-time patterns across a run predict fluctuations of the BOLD signal during experimental blocks. In addition to having implications for studies investigating individual differences in response times and SD_BOLD_, these findings are also relevant for studies investigating differences in SD_BOLD_ between experimental conditions. For example, Garrett et al. (2014) showed widespread reductions in SD_BOLD_ as task difficulty in a face-matching task increased, and that the extent of SD_BOLD_ reduction across difficulty levels predicted increases in response time and reductions in accuracy. These patterns of results were interpreted as resulting from the brain’s dynamic range or edge of criticality being modulated by task difficulty, such that more difficult tasks constrain the range of states brains can occupy. However, when all task difficulty levels consist of the same trial length, an alternative explanation is simply that as RTs increase with task difficulty, SD_BOLD_ necessarily reduces due to the reduction in on-task/off-task toggling described earlier. Similarly, Grady and Garrett (2018) showed that externally directed cognitive tasks induce increased variability relative to internally directed tasks in young (but not older) adults, which was interpreted as BOLD variability modulating in response to changes in environmental uncertainty—an effect that diminishes with age. However, examination of the task design and response-time patterns allow for an alternative explanation. In block designs with fixed trial lengths (3 seconds in this case), BOLD variability should systematically decrease as response times increase. Indeed, in young adults, the response time patterns showed that externally directed tasks showed faster response-times (and increased BOLD variability) compared to internally directed tasks. In addition, this pattern was not observed in older adults (i.e., there were no differences in response-times or BOLD variability between internally and externally directed tasks).

Furthermore, while the current study does not directly address the relationship between aging, cognitive performance and BOLD variability, the findings are directly relevant to such studies. First, it should be noted that earlier studies investigating this relationship failed to tease apart the independent effects of age and cognitive performance on BOLD variability (Garrett, Kovacevic, et al., 2011, 2013), which is potentially problematic as age and response-times are themselves related. That is, it could be the case that some proportion of the age-differences in BOLD variability in these studies is attributable to differences in response times between young and older adults, and that these response-time differences—according to the framework proposed here—drives the differences in BOLD variability between the age groups. Interestingly, in direct opposition to most of the literature, Boylan et al. (2021) have shown *greater* BOLD variability to be associated with increased age and reduced cognitive performance. Similarly, in an episodic memory study, Wang et al. (2022)showed that BOLD variability in a widespread set of regions *increased* with age in female participants, albeit with more mixed findings in males: some regions showed age-related increases and others showed decreases. We tentatively propose these findings may be due to both studies *not* using a design that induced a toggling between on and off-task during experimental conditions: Boylan et al. (2021) used a block-design of an N-back paradigm in which participants are continuously on-task during the run due to constantly having to maintain the item presented N items previously; Wang et al. (2022) used an event-related design in which only the HRF function for each trial was analysed.

To summarise, we have presented findings that undermine two key claims made in the literature regarding BOLD variability. First, the most common method of computing task-related SD_BOLD_ is not independent of mean_BOLD_: knowing the mean percent signal change of a set of voxels allows reasonable prediction of their SD_BOLD_ values. To mitigate the effects of mean on SD analyses in future studies, we propose regressing out the experimental design prior to calculating SDBOLD. Interestingly, in a study that used machine-learning algorithms to investigate how well BOLD variability was able to predict participant identity and cognitive task, regressing out the task design increased participant prediction but decreased task prediction (Gaut et al., 2019). One interpretation of the latter effect is that, prior to regressing out the experimental design, task-specific mean_BOLD_ effects contributed to BOLD variability spatial patterns thereby increasing the ability of SD_BOLD_ to predict cognitive tasks. Second, across a number of analyses, we have provided evidence that switching between on- and off-task states in blocked designs with fixed trial-lengths contributes to BOLD variability and underlies the relationship between SD_BOLD_ and response times. To mitigate the effects of this task-switching, we propose that researchers adopt designs that involve self-paced trials within an experimental block. Adopting this procedure would result in a purer measure of BOLD variability that indexes fluctuations of the BOLD signal while engaged in a task as opposed to a measure that is heavily influenced by the switching between cognitive states.

In conclusion, interest in brain signal variability and brain dynamics more generally has undergone a tremendous surge in the last decade and has provided important insights into the nature of brain function. The results of the present study, however, indicate that current measures of task-related BOLD variability are sensitive to aspects of the experimental design and its interaction with the specifics of how BOLD variability is calculated. A promising avenue for further research would be using large open-access datasets—ideally involving multiple research groups—to clarify how different experimental designs and analytic pipelines affect the outcomes of specific research questions.

## Footnotes

1 Note that the pre-registration of the current study specified that additional noise components would be classified manually. However, visual inspection of the ICs classified by AROMA as noise components indicated that physiological and scanner artifacts were picked up by AROMA, and so manual classification was deemed to not be necessary.

